# Identification of loop regions as motifs determining cellular and organ chirality in Myosin 1C

**DOI:** 10.64898/2026.04.14.718603

**Authors:** Asuka Yamaguchi, Takeshi Sasamura, Kohei Yoshimura, Takeshi Haraguchi, Toshifumi Mori, Kohji Ito, Kenji Matsuno

## Abstract

Left–right (LR) asymmetry occurs throughout the animal kingdom, from microscopic to macroscopic scales; however, how microscale LR asymmetry (chirality) is integrated into higher-order structural LR asymmetry remains unclear. In *Drosophila*, the actin motor proteins Myosin1D and Myosin1C impose opposite chirality states on cell shape and intracellular F-actin flow, thereby directing dextral and sinistral morphogenesis, respectively. However, the protein motifs that confer their distinct activities in specifying chiral states have remained unknown. Here, we show that loop motifs within the myosin head domain that form actin-binding sites determine enantiomeric chirality. By swapping these loop motifs between Myosin1D and Myosin1C, we demonstrate that Myosin1D acquires sinistral activity when carrying Myosin1C loops. AlphaFold3 and molecular dynamics analyses reveal differences in loop structure and actin-binding energetics that may account for the contrasting chiral activities of Myosin1D and Myosin1C. Our findings identify specific protein motifs that dictate the handedness of actin flow and organ asymmetry, providing a mechanistic link between molecular interactions and macroscopic left–right patterning.

## Introduction

Chirality is a fundamental property observed at the molecular, cellular, and organismal levels. A three-dimensional object is defined as chiral if it does not superimpose on its mirror image, a form of left-right (LR) asymmetry. At the molecular level, numerous biomolecules, including amino acids and sugars, exhibit homochirality — an essential prerequisite for biological order. Organisms rely on this uniform chirality, and concerns have been raised that mirror-image life forms could pose risks such as immune evasion(*1*).

Beyond molecules, many organs and body structures exhibit intrinsic chirality that underlies their physiological and behavioral functions(*2*). For instance, the looping of the vertebrate heart is essential for correct blood flow(*3, 4*), while snail shells coil chiral(*5*), and their predators have evolved jaws adapted to this LR asymmetry(*6*).

Whether such organ-level chirality originates from molecular chirality remains unresolved. The “F-molecule” model proposes that a molecular chiral determinant governs organ and body asymmetry(*7*). In mice, the basal body of cilia is considered an F-molecule: clockwise ciliary rotation generates a leftward flow of the extra-embryonic fluid in the node, which activates asymmetric gene expression(*8–11*). However, LR asymmetry in invertebrates and certain vertebrates, including birds and frogs, develops independently of cilia and nodal flow(*12*). Therefore, further research is required to gain a comprehensive understanding of the molecular-level origins of organ chirality and how molecular chirality determines organ chirality.

At the cellular level, eukaryotic cells, which serve as an intermediate level connecting molecular processes and organ-level organization, display inherent chirality in morphology(*13–16*), motility(*17–19*), and intracellular dynamics(*20–23*). This phenomenon, termed “cell chirality,” occurs in both cultured cells and tissues such as the *Drosophila* hindgut and genitalia, the embryonic chicken heart(*24*), and mouse vasculature(*15*). Cell chirality largely depends on the cytoskeletal network, particularly actin filaments and associated factors such as formin(*25*), fascin(*26*), and non-muscle myosin II(*22, 27*). These factors also contribute to LR-asymmetric morphogenesis in *Drosophila*(*28, 29*), snails(*30*), and *C*. *elegans*(*31*). Yet, how cytoskeletal chirality is formed and amplified into tissue- and organ-level asymmetry remains largely unclear.

In *Drosophila*, two class I myosins, *Myosin1D* (*Myo1D*) and *Myosin1C* (*Myo1C*), determine right-handed (dextral) and left-handed (sinistral) chirality, respectively(*32–35*). *Myo1D* directs right-handed morphogenesis of the embryonic gut and genitalia, and its loss reverses LR polarity, resulting in left-handed chirality (*19, 32, 33, 36*). Conversely, ectopic expression of *Myo1C* induces mirrored organ and cell chirality, leading to left-handed chiralty(*32, 34, 35*). Both myosins can even generate chirality *de novo* in other bilateral tissues(*35*).

In *Drosophila* macrophages overexpressing *Myo1D* or *Myo1C*, cytoplasmic F-actin flows clockwise and counterclockwise, respectively^37^. An *in vitro* motility assay further revealed that surface-immobilized Myo1D drives chiral movement of single F-actin filaments on the cover glass(*35, 37*). When actin concentration increases, Myo1D organizes F-actin bundles into ring-like structures, termed actin chiral rings, which rotate clockwise^37^. This rotation suggests that Myo1D assembles F-actin into a circular arrangement with barbed-end-to-pointed-end polarity aligned in a specific direction. These findings suggest that the Myo1D–actin interaction represents the molecular origin of cell chirality. Studies have begun testing this hypothesis, and progress is being made in biophysical studies of Myo1D and in the structural analysis of its mammalian paralog, which also induces chiral F-actin movement(*37–39*). However, the precise mechanism by which Myo1D induces chiral actin movement, and the specific structural domains of Myo1D required for this activity, remain unresolved. Moreover, Myo1C fails to generate chiral actin flow in *vitro*, instead producing random motion(*35, 37, 40*). Therefore, the molecular basis of the left-handed chirality mediated by Myo1C is still unknown.

*Drosophila* Myo1D and Myo1C are evolutionarily conserved proteins composed of three domains: a head, a neck (IQ), and a tail domain(*41*). Previous studies identified the head domains of Myo1D and Myo1C as responsible for right- and left-handed chirality, respectively, in organs and cells(*35, 42, 43*). The head domain contains actin-interacting regions that generate movement; thus, the force-bearing chirality of F-actin is likely determined within this region(*44*). Myosins bind actin through several surface loops—loop 2, loop 3, loop 4, cm loop, helix–turn–helix loop, and activation loop(*44, 45*). These loops vary considerably among myosin families and contribute to their distinct biochemical activities(*44*). For example, previous studies have shown that exchanging these loops between myosin species with distinct ATPase activities transfers their enzymatic properties, indicating that these loops encode the functional specificity of myosin(*46*).

We hypothesized that differences in these actin-interacting loops define the right- and left-handed chirality of Myo1D and Myo1C. Although the overall head domains of Myo1D and Myo1C share sequence similarity(*41*) (∼50%), their loop regions diverge markedly (Fig. 1A). We therefore predicted that variations in loop composition underlie their opposing chiral activities. To test this, we engineered chimeric myosin genes in which individual loops were swapped between Myo1D and Myo1C (Fig. 1B). These chimeric constructs were overexpressed in the *Drosophila* embryonic gut to determine which loops are sufficient for left- and right-handed activities. Additionally, to examine whether loop swapping alters left- or right-handed states of cellular chirality, we analyzed F-actin flow in macrophages expressing these chimeric myosins. We found that swapping loops of Myo1D to those of Myo1C in the backbone of Myo1D converted its activity from right- to left-handed in both gut morphology and macrophages’ F-actin flow. Conversely, introducing Myo1D loops into Myo1C did not reverse chirality but reduced its left-handed activity. Furthermore, structural modeling of Myo1D-actin or Myo1C-actin complexes using AlphaFold3(*47*), together with molecular dynamics (MD) simulations, reveals that the Myo1C loops interact with actin in a similar manner both when it is in the Myo1C backbone and when it is transferred to the Myo1D backbone. Based on these results, we propose that the cellular and organ chirality of Myo1C arises from loop structures in the myosin head.

**Fig. 1.**
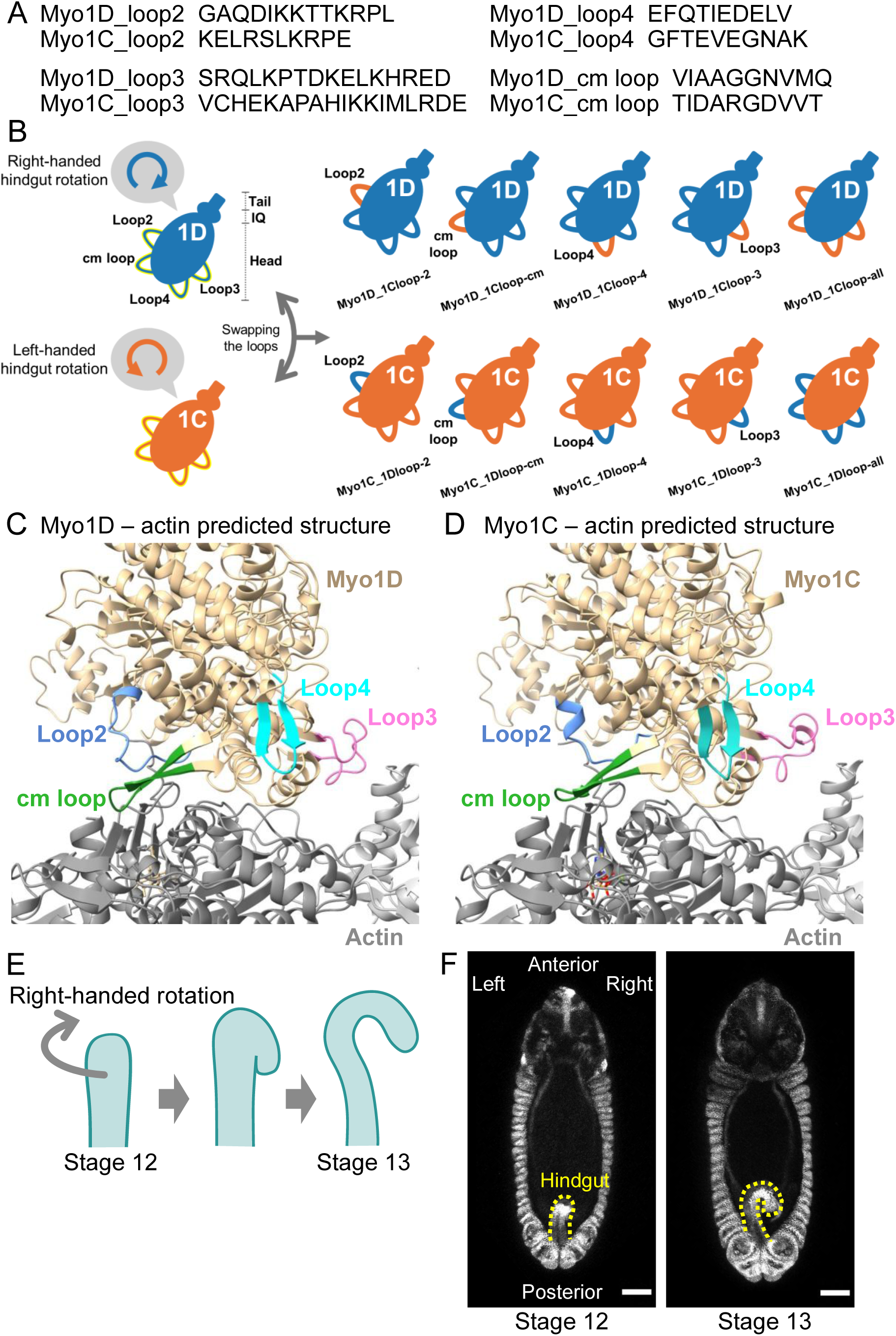
Scheme of the loop-swapping experiment. **(A)** Amino acid sequences of each loop of Myo1D and Myo1C. **(B)** Diagram of myosin structures and a scheme of the loop-swapping experiment. Orange diagram; Myo1C, blue diagram; Myo1D. The head domain, IQ domain, and tail domain are indicated next to the Myo1D diagram, and they can be applied to the Myo1C diagram. Loop2, cm loop, loop3, and loop4 are shown in the Myo1D diagram and can be applied to the Myo1C diagram. Clockwise and Counterclockwise directions of round arrows represent the right- and left-handed rotation of the embryonic gut, respectively, induced by their overexpression. **(C, D)** Predicted structures of Myo1D-actin complex and Myo1C-actin complex by AlphaFold3. The structure of Myo1D (C) and Myo1C (D); ocher, actin monomers; gray, loop2; blue, cm loop; green, loop4; cyan, loop3; pink. **(E)** Diagrams of hindgut morphogenesis. **(F)** The hindgut (shown by a yellow dashed line) in a wild-type embryo at stages 12 (left panel) and 13 (right panel).

## Results

### Swapping loops alters right- and left-handed activities of *Myo1D* and *Myo1C*, respectively, in the hindgut

Previously, high-resolution cryo-EM structures of mouse class 1 myosin revealed the actin-binding states through the following loops: loop2, loop3, loop4, and the cm loop(*48*). These loops are located at the actin-binding surface of *Drosophila* Myo1D and Myo1C according to the predicted structures of Myo1D- and Myo1C-actin complex by AlphaFold3(*47*) (Figures 1C and 1D). Each loop region consists of approximately 10-18 amino acid residues, and the amino acid sequences of the corresponding loops in Myo1D and Myo1C differ markedly. (Figures 1A, 1C and 1D). To test the possibility that these loop regions determine the right- and left-handed activities of Myo1D and Myo1C, respectively, we created the chimeric myosin genes by swapping the regions encoding each loop between the cDNAs of *Myo1D* and *Myo1C* (Figure 1B).

Roles of *Myo1D* and *Myo1C* in the formation of organ chirality that arises from cell chirality have been studied in the embryonic hindgut and the male genital disk(*13, 28, 33, 42*). In wild-type embryos, the hindgut initially forms as a bilaterally symmetrical structure with its anterior end flexed ventrally (Figures 1E and 1F). Then it starts to rotate anticlockwise as viewed from the posterior end at stage 12(*36, 49*) (Figures 1E, 1F and Supplementary Movie 1). The hindgut torsion stops after twisting 90 degrees at stage 13, resulting in the right-curved structure of the hindgut at 100 % as viewed from the dorsal side of the embryo, designated as right-handed(*32, 36, 49*) (Supplementary Movie 1, Figures 1E, 1F, 2A and 2F - row①). On the other hand, embryos homozygous for the *Myo1D* null mutant, *Myo1D^L152^*, showed LR-inversion of hindgut rotation at around 75%, designated as left-handed, as reported previously(*32*) (Figures 2B and 2F- row①). Using the Gal4/UAS system, misexpression of wild-type *Myo1D* driven by a hindgut-specific Gal4 driver, *byn* (*brachyenteron*)*-gal4*(*50*), completely rescued the LR-inversion in these mutant embryos, as *Myo1D* has the right-handed activity(*32, 34*)(Figure 2C and 2F - row②). Using the same system, the left-handed activity of *Myo1C* was detected by overexpressing wild-type *Myo1C*, as the LR inversion of hindgut rotation was induced in wild-type and enhanced from 75 to 100% in *Myo1D^L152^* homozygous embryos(*32*) (Figure 2D and 2F - row③). Based on these LR phenotypes of the embryonic hindgut, we here evaluated the right- and left-handed activities of *Myo1D* and *Myo1C* derivatives, which have their loop regions swapped alternately (Figures 1B and 2F).

**Fig. 2.**
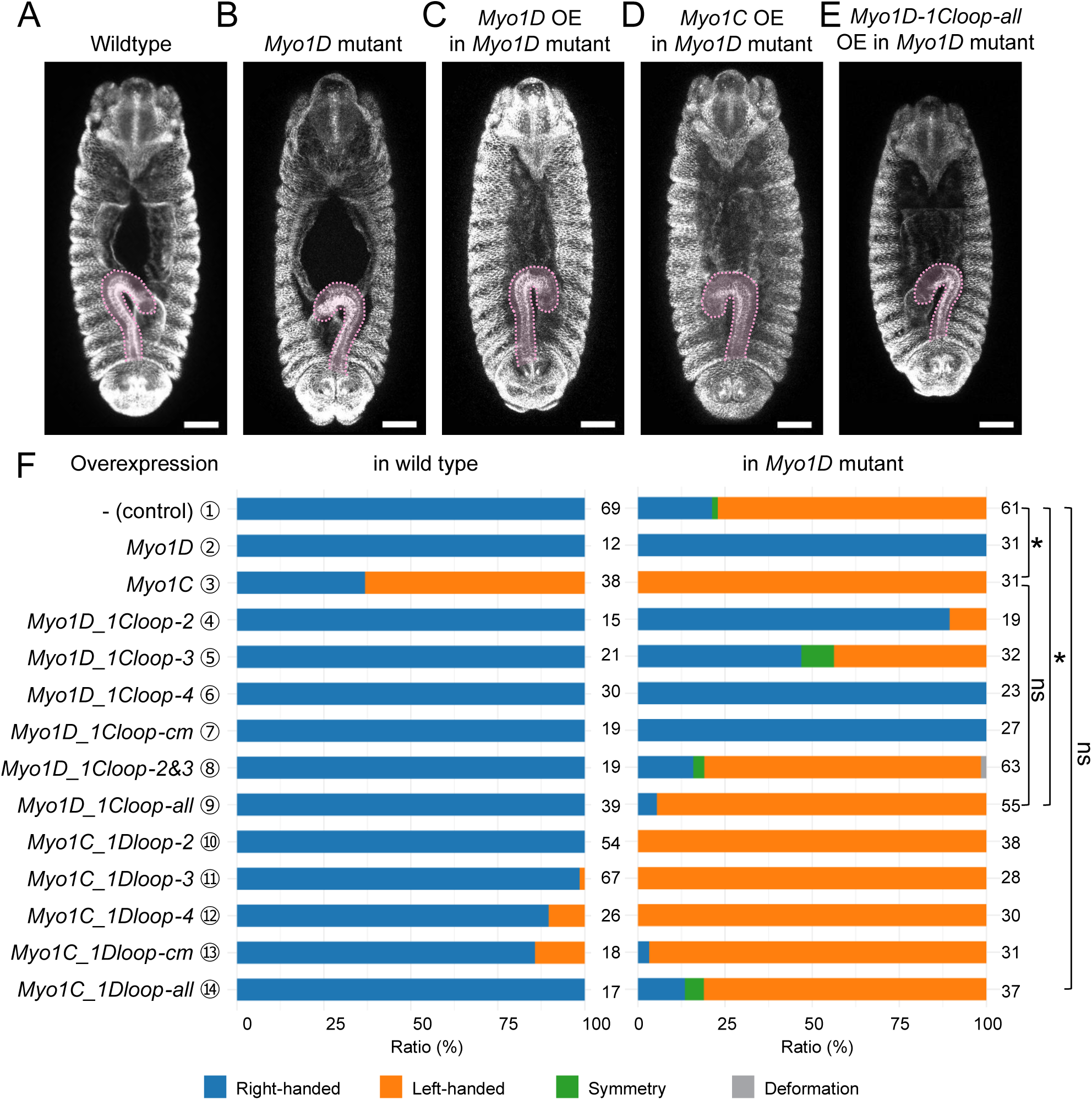
Loop swapping alters the activity of Myo1D and Myo1C for the LR asymmetric hindgut rotation. (A, **B)** Representative phenotype of the hindgut at stage 13 in *w^1118^*homozygote as wild type (A) and *Myo1D^L152^* homozygote (B). **(C-E)** Representative phenotypes of hindgut in stage 13 of *Myo1D^L152^*homozygote overexpressing (indicated by OE) wild-type *Myo1D* (C), wild-type *Myo1C* (D), or *Myo1D-1C all loops* (E). In A-E, embryos were stained with a FasIII antibody (white), and the hindgut was shown in light pink. **(F)** The bar graphs show the frequency (%) of phenotypes associated with LR-asymmetric hindgut rotation. In bar graphs, blue, orange, green, and gray represent right-handed rotation, left-handed rotation, LR symmetry, and deformation, respectively, as shown in the bottom. The left bar graphs show the frequency of the phenotypes in wild-type embryos (-(control)) and in embryos overexpressing the respective genes, indicated at the left. The right bar graphs show the frequency of the phenotypes in *Myo1D^L152^* homozygote embryos (-(control)), and embryos overexpressing the respective genes, indicated at the left. The number of embryos observed is shown on the right side of each bar. The numbers enclosed in circles correspond to the structures of the chimeric myosins obtained by loop swapping between Myo1D and Myo1C. The *P*-values were obtained by Fisher’s exact test with Holm correction; ns and * represent *P* > 0.05 and *P* < 0.05, respectively.

These derivatives are designated as *Myo1D_Myo1C”loop name”* for the *Myo1D* backbone carrying *Myo1C* loops, and *Myo1C_Myo1D”loop name”* for the *Myo1C* backbone carrying *Myo1D* loops (Figure 1B). We first analyzed the specific activity of each loop in Myo1D and Myo1C for the hindgut’s right- and left-handed chirality. *Myo1D_1Cloop-2*, *Myo1D_1Cloop-3*, *Myo1D_1Cloop-4,* and *Myo1D_1Cloop-cm* are swapped chimeric *Myo1D* genes in which the sequences encoding loop2, loop3, loop4, and cm loop were respectively swapped with the corresponding loops of Myo1C in the backbone of Myo1D (Figure 1B).

*Myo1C_1Dloop-2*, *Myo1C_1Dloop-3*, *Myo1C_1Dloop-4*, and *Myo1C_1Dloop-cm* are swapped chimeric Myo1C genes in which the sequences encoding loop2, loop3, loop4, and cm loop were respectively swapped with the corresponding loops of Myo1D in the Myo1C background (Figure 1B). To confirm whether the swapped *Myo1D* and *Myo1C* genes produce the predicted derivative proteins, we conducted Western blot analysis (Figures S1 and S2). As these derivatives have mGFP- or mCherry-tags at their C-termini, we specifically overexpressed them in the embryonic hindgut driven by *byn-Gal4* and detected by using anti-mGFP or anti-mCherry antibody. We detected wild-type Myo1D and Myo1C, and all of their derivatives, at the predicted sizes (approximately 144 kDa) (Figures S1 and S2).

As described before, the wild-type hindgut (control) showed 100 % right-handed rotation (Figure 2A and 2F - row①). Overexpression of wild-type *Myo1D* in the hindgut of wild-type embryos driven by *byn-Gal4* showed 100 % right-handed rotation similar to that of control (Figure 2F - row②). However, overexpression of wild-type *Myo1C* in the hindgut of wild-type embryos resulted in 62% left-handed rotation, as the wild-type Myo1C has left-handed activity (Figure 2F - row③). We found that overexpression of the chimeric Myo1D genes, *Myo1D_1Cloop-2*, *Myo1D_1Cloop-3*, *Myo1D_1Cloop-4*, and *Myo1D_1Cloop-cm*, resulted in right-handed rotation of the hindgut (Figure 2F - row④ to ⑦). These results suggested that either chimeric *Myo1D* genes maintained or lost right-handed activity but did not gain left-handed activity in the wild-type hindgut. We also examined the phenotypes of overexpression of the chimeric *Myo1C* genes, *Myo1C_1Dloop-2*, *Myo1C_1Dloop-3*, *Myo1C_1Dloop-4,* and *Myo1C_1Dloop-cm*. We reveal that overexpression of *Myo1C_1Dloop-4* or *Myo1C_1Dloop-cm* induced left-handed rotation at around 12.5 %, which was less than that of wild-type *Myo1C* (62%), demonstrating that the left-handed activity of them was maintained but reduced (Figure 2F - row ⑫ and ⑬). We also found that overexpression of *Myo1C_1Dloop-2* or *Myo1C_1Dloop-3* in the hindgut hardly induced left-handed rotation, suggesting that their left-handed activity was lost (Figure 2F - row ⑩ and ⑪). However, when right- and left-handed activities were lost or reduced in our experiments, the possibility of non-specific inactivation via protein denaturation or misfolding caused by loop swapping cannot be ruled out.

We also evaluated the right- and left-handed activities of chimeric Myo1D and Myo1C in the *Myo1D* mutant. To that end, we overexpressed the wild-type *Myo1D*, wild-type *Myo1C*, chimeric *Myo1D*, and chimeric *Myo1C* genes in the embryonic hindgut of *Myo1D^L152^* homozygote driven by *byn-Gal4*. The hindgut of *Myo1D^L152^* homozygote showed 75 % left-handed rotation, as reported before(*32, 43*) (Figures 2B and 2F - row①). Overexpression of wild-type *Myo1D* in *Myo1D^L152^* homozygote resulted in 100% right-handed rotation, demonstrating that the left-handed state was suppressed entirely(*32, 43*) (Figures 2C and 2F - row②). Significantly, overexpression of wild-type *Myo1C* in the hindgut of *Myo1D^L152^* homozygote increased the left-handed rotation from 75% to 100 %, demonstrating that we can detect the left-handed activity of *Myo1C* in this system (Figure 2D and 2F - row③). Since *Myo1D^L152^* is a null mutant of *Myo1D*, these results suggest that the left-handed activity of *Myo1C* is independent of *Myo1D*, as reported before(*34*). The activities of these wild-type *Myo1D* and *Myo1C* were compared with those of chimeric *Myo1D* and *Myo1C* to evaluate the potential activities of each loop region (Figure 1B). We found that the hindgut overexpressing *Myo1D_1Cloop-2* or *Myo1D_1Cloop-3* still shows left-handed rotation at 10% and 43%, respectively (Figure 2F - row④ and ⑤), whereas overexpression of wild-type *Myo1D* completely suppresses it (Figure 2F - row②). Therefore, the right-handed activity of *Myo1D_1Cloop-2* and *Myo1D_1Cloop-3* are reduced, although the reduction can be attributed to protein denaturation or misfolding *via* the swapping of loops.

Conversely, overexpression of *Myo1D_1Cloop-4* or *Myo1D_1Cloop-cm* led to 100 % right-handed rotation, which was the same as the consequence of wild-type *Myo1D* overexpression (Figure 2F - row⑥and ⑦). Therefore, loop4 and cm loop of Myo1C function within the backbone of Myo1D in the same manner as those of Myo1D. Given that Myo1C_1Dloop-4 or Myo1C_1Dloop-cm maintained the left-handed activity, although it was reduced, upon the overexpression in wild-type hindgut (Figure 2F - row ⑫ and ⑬), these results suggest that the loop4 and the cm loop are exchangeable between Myo1D and Myo1C for their respective activity of right- and left-handed rotations associated with their backbones.

Using the same approach, we also analyzed the specific requirement of the loop regions in Myo1C for its left-handed activity in *Myo1D^L152^* homozygote. Given that overexpression of wild-type *Myo1C* in the hindgut of *Myo1D^L152^* homozygote increased the frequency of left-handed rotation from 75 to 100 %, we examined whether this activity is maintained in *Myo1C_1Dloop-2*, *Myo1C_1Dloop-3*, *Myo1C_1Dloop-4,* and *Myo1C_1Dloop-cm* in the hindgut of *Myo1D^L152^* homozygote. We found that overexpression of *Myo1C_1Dloop-2*, *Myo1C_1Dloop-3*, *Myo1C_1Dloop-4*, or *Myo1C_1Dloop-cm* induced almost 100 % left-handed rotation (Figure 2F - row⑩ to ⑬). Therefore, swapping of any one of the loop regions in Myo1C to the corresponding loop regions of Myo1D did not affect the left-handed activity of Myo1C. As described above, in the wild-type hindgut, *Myo1C_1Dloop-2* and *Myo1C_1Dloop-3* showed a decreased left-handed activity, as compared with wild-type *Myo1C* (Figure 2F - row③, ⑩ and ⑪). This might be explained if endogenous *Myo1D* counteracts the weakened left-handed rotation induced by those chimeric *Myo1C* in wild type, as such counter-activity is absent in the hindgut of *Myo1D^L152^* homozygote.

We speculate that these loop regions may act together to exhibit right- and left-handed activities. Specifically, regarding loop2 and loop3 of Myo1D, individual replacement of loop 2 or loop 3 of Myo1D with the corresponding Myo1C loop led to reduced right-handed activity of Myo1D in the *Myo1D* mutant (Figure 2F - row④ and ⑤). This implies that these two loops of Myo1D may exhibit a synergistic activity to induce the right-handed rotation of the hindgut. To test this idea, we analyzed a *Myo1D* gene variant where loop2 and loop3 were swapped with the corresponding regions from Myo1C within the Myo1D backbone (Myo1D_1Cloop-2&3). Extending this further, we analyzed the activities of Myo1D-Myo1C chimeras generated by simultaneously exchanging all four loop regions between the two myosins. (*Myo1D_1Cloop-all* and *Myo1C_1Dloop-all*) (Figure 1B). Those chimeric Myo1D and Myo1C proteins overexpressed in the hindgut were detected at the predicted molecular weight (144 kDa) by Western blot analysis. (Figure S1 and S2).

We found that overexpression of the *Myo1D_1Cloop-2&3* gene in the hindgut of wild-type embryos, driven by *byn-Gal4*, resulted in 100% right-handed rotation (Figure 2F - row⑧). Thus, *Myo1D_1Cloop-2&3* did not demonstrate left-handed activity in wild type, although we cannot determine whether it maintains right-handed activity or loses it. We also overexpressed *Myo1D_1Cloop-2&3* in the hindgut of *Myo1D^L152^* homozygote, resulting in 79% left-handed rotation, which was similar to the phenotype observed in *Myo1D^L152^* homozygote without the overexpression (Figure 2F - row⑧). These results suggest that *Myo1D_1Cloop-2&3* completely lost right-handed activity. Considering that overexpression of *Myo1D_1Cloop-2* or *Myo1D_1Cloop-3* in *Myo1D^L152^* homozygote resulted in the partial reduction of the left-handed rotation (Figure 2F - row ④ and ⑤), our results suggest that the loop 2 and the loop 3 regions of Myo1D are involved in the right-handed activity in a coordinated manner, although we do not exclude the possibility that the activity of Myo1D_1Cloop-2&3 was reduced due to protein denaturation or misfolding.

We then analyzed the activity of *Myo1D_1Cloop-all*. In the wild-type hindgut, overexpression of *Myo1D_1Cloop-all* resulted in 100% right-handed rotation (Figure 2F - row⑨), demonstrating that *Myo1D_1Cloop-all* did not show left-handed activity. However, we revealed that overexpression of the *Myo1D_1Cloop-all* gene in the hindgut of *Myo1D^L152^* homozygote increased the left-handed rotation from 75 to 95 % (Figure 2F - row⑨), which was similar to the effect of wild-type *Myo1C* overexpression (100% left-handed rotation) in *Myo1D^L152^* homozygote (Figure 2F - row③). These results suggest that *Myo1D_1Cloop-all* acquired left-handed activity by swapping the four, but not one or two, loop regions from Myo1D to Myo1C in the Myo1D backbone. We checked the localization of the Myo1D_1Cloop-all protein in epithelial cells of the hindgut and could not find any detectable differences compared to the localization of wild-type Myo1D and Myo1C (Figure S3), indicating that the intracellular localization is not involved in the acquired left-handed activity of the Myo1D_1Cloop-all protein. Based on these results, we hypothesized that the four loop regions of Myo1C contain at least some of the information necessary for left-handed activity.

Although we found that *Myo1D_1Cloop-all* showed left-handed activity in *Myo1D^L152^* homozygote, overexpression of *Myo1D_1Cloop-all* in the wild-type hindgut did not induce left-handed rotation at all (Figure 2F - row⑨). We predicted that this seemingly contradictory result is because, in the wild-type hindgut, the right-handed rotation induced by endogenous *Myo1D* counteracts the relatively weak left-handed rotation induced by *Myo1D_1Cloop-all*. Considering this possibility, we checked the effect of *Myo1D_1Cloop-all* and wild-type *Myo1C* in the hindgut of embryos heterozygous for *Myo1D^L152^* in which only one wild-type *Myo1D* gene, instead of two, exists. As the LR-phenotype of the hindgut in *Myo1D^L152^*heterozygote may be more sensitive to genetic background than that of the wild type, we overexpressed *Lifeact-mGFP6* in the hindgut of *Myo1D^L152^*heterozygote driven by *byn-Gal4* as a control. We found that left-handed rotation was not induced (Figure 3). However, overexpression of wild-type *Myo1C* induced 100 % left-handed rotation under the same conditions (Figure 3). As previously mentioned, when wild-type *Myo1C* was overexpressed in wild-type embryos, the incidence of left-handed rotation was 62% (Figure 2F -row③). Therefore, these results indicate that in wild-type embryos, the left-handed rotation induced by overexpression of *Myo1C* is suppressed by the right-handed rotation induced by endogenous *Myo1D*. As we predicted, in the hindgut of *Myo1D^L152^* heterozygote, overexpression of *Myo1D_1Cloop-all* induced 10 % of left-handed rotation, demonstrating that *Myo1D_1Cloop-all* has a left-handed activity (Figure 3). We speculate that this left-handed activity of *Myo1D_1Cloop-all* is masked and undetectable in the presence of two copies of the wild-type *Myo1D* gene. According to these results, we concluded that the four loop regions of Myo1C together contain the information necessary for left-handed activity. We also examined the effect of *Myo1D_1Cloop-2&3* in the hindgut of embryos heterozygous for *Myo1D^L152^*.

**Fig. 3.**
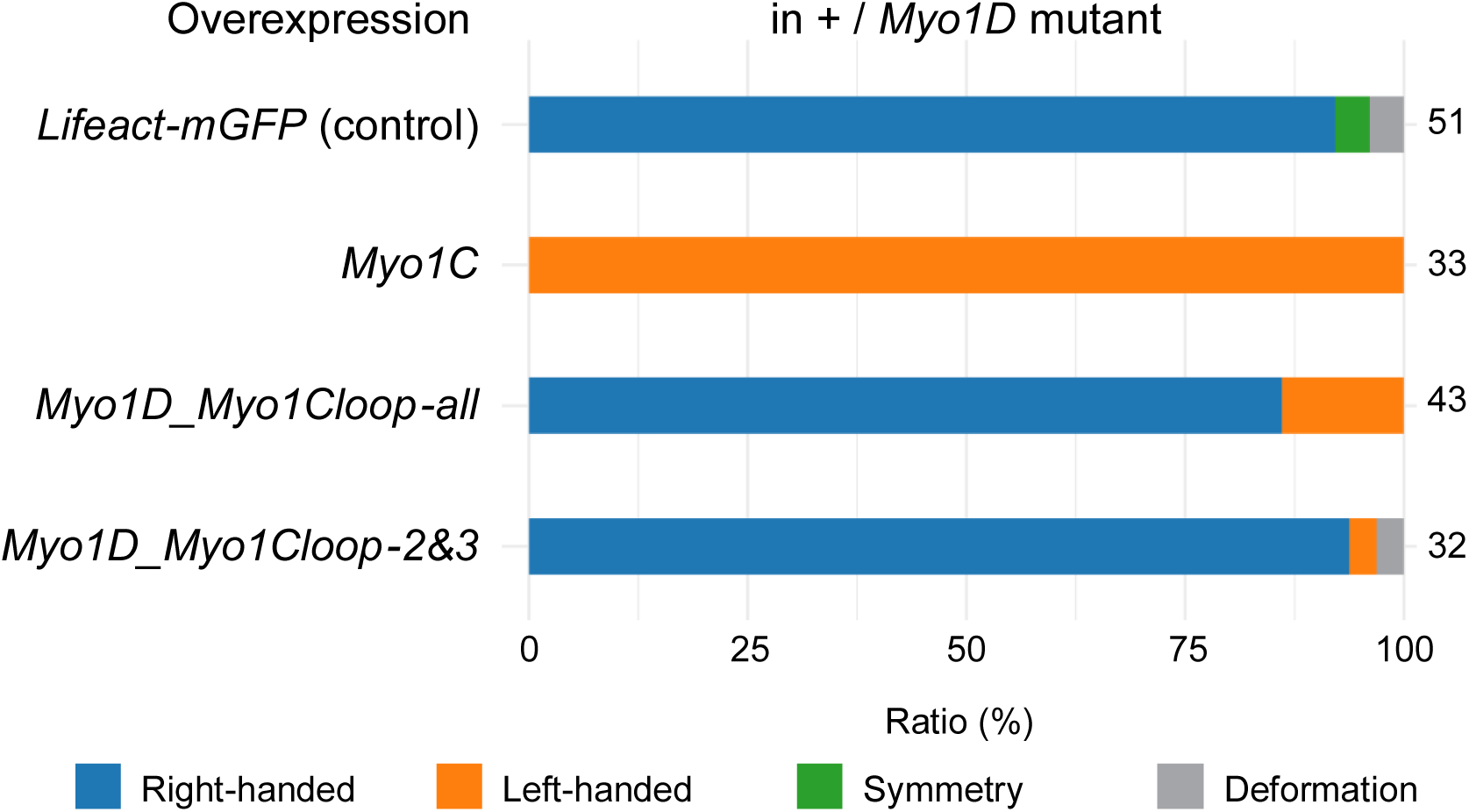
Myo1D, where all four loops are replaced with those of Myo1C, induces left-handed hindgut rotation in *Myo1D* heterozygous embryos. The bar graphs show the frequency (%) of phenotypes associated with LR-asymmetric hindgut rotation (indicated at the bottom) in *Myo1D^L152^* heterozygotes overexpressing the respective genes indicated at the left. The number of embryos observed is shown on the right side of each bar. In bar graphs, blue, orange, green, and gray represent right-handed rotation, left-handed rotation, LR symmetry, and deformation, respectively, as shown at the bottom.

Overexpression of *Myo1D_1Cloop-2&3* rarely induced the LR inversion (Figure 3), indicating that left-handed activity cannot be conferred by substitution of only loop 2 and loop 3 from Myo1C into the Myo1D backbone, but instead requires exchange of all four loops (Figure 2F -row⑨ and Figure 3). This result is consistent with the fact that overexpression of *Myo1D_1Cloop-2&3* did not affect the LR-phenotypes in *Myo1D^L152^*homozygous embryos (Figure 2F -row⑧).

We next explore the possibility that all four loops of Myo1D function in coordination to determine its right-handed activity. We found that overexpression of *Myo1C_1Dloop-all* resulted in 80 % left-handed rotation in *Myo1D^L152^* homozygote (Figure 2F - row⑭), which is similar to the frequency of left-handed rotation in *Myo1D^L152^* homozygote (Figure 2F - row①). Therefore, we did not detect right- or left-handed activity of *Myo1C_1Dloop-all*. These results suggest that while the four loops of Myo1C transferred to the backbone of Myo1D retain their original left-handed activity (Figure 2F -row⑨), the four loops of Myo1D lack such capability.

### The four loop regions of Myo1C have an activity dictating the counterclockwise rotation of F-actin flow in *Drosophila* macrophages

Our analysis using the swapped *Myo1D* and *Myo1C* genes *in vivo* revealed that the four loop regions of Myo1C, together, comprise sufficient information to impose left-handed activity on the Myo1D background. Given that the loop regions are known to interact with F-actin(*44*), we next investigated the effects of Myo1D and Myo1C, in which the four loop regions were simultaneously swapped, on the cytoplasmic dynamics of F- actin, with a specific focus on its chiral dynamics. We have reported an experimental system to quantitatively evaluate the chirality of F-actin flow in the cytoplasm of *Drosophila* macrophages(*51*). In this system, we collect the macrophages from the body fluid of third-instar larvae, culture them *ex vivo*, and analyze the chirality of F-actin flow *via* time-lapse imaging(*51, 52*) (Figure 4A). Time-lapse movies of these macrophages overexpressing *Lifeact-mCherry* driven by macrophage-specific *Gal4* driver, *He-Gal4*, showed a typical retrograde flow of F-actin and allowed us to observe intracellular dynamics of F-actin (Figure 4B and Supplementary Movie 2). As described before(*51*), we obtained the directional vector for each pixel predicted by optical flow estimation at the respective time points of the time-lapse images for 10 min(*53*) (Figure 4C). Subsequently, the directional vector was decomposed into two vectors: the unit vector (*a⃗_i_*) in the direction of a cell centroid and the vector that is perpendicular to the unit vector (*C⃗_i_*) (Figure 4C). The latter vector, called the “chiral vector”, represents the chiral (rotational) movement of each pixel (Figure 4C). If the chiral vector indicates clockwise (right-handed) and counterclockwise (left-handed), chiral vectors were marked by negative (-) and positive (+) values, respectively (Figure 4C). To quantitatively evaluate the chirality of F-actin in each experimental condition, we summed all values of the chiral vectors in each cell, and these sums were obtained as the “chiral index”. Consistent with a previous study(*51*), wild-type macrophages (control) showed a clockwise bias in the chiral index, and this bias was significantly enhanced by overexpression of wild-type *Myo1D*, demonstrating the right-handed activity of *Myo1D* (Figure 4D and Table 1). However, as previously reported, overexpression of wild-type Myo1C induced a significant counterclockwise bias (Figure 4D and Table 1). Therefore, wild-type *Myo1D* and *Myo1C* have right- and left-handed activities, respectively, in the circular flow of F-actin, consistent with their corresponding activities in the hindgut as described above. Using this system, we investigated the effects of swapping the loop regions between the Myo1D and Myo1C on the chirality of F-actin circular flow. We revealed that overexpression of *Myo1D_1Cloop-all* induced significant counterclockwise bias in the chiral index, same as overexpression of wild-type *Myo1C* (Figure 4D and Table1). This indicates that the swapping of all four loop regions of Myo1D to those of Myo1C in the Myo1D backbone alters its activity from right-handed to left-handed, not only in the hindgut rotation, but also in circular F-actin flow in macrophages. On the other hand, overexpression of *Myo1C_1Dloop-all* showed no bias in the chiral index (Figure 4D and Table1). Considering that wild-type *Myo1C* overexpression induced significant biases to left-handed F-actin rotation, we speculated that the left-handed activity of Myo1C_1Dloop-all decreases as compared with that of wild-type *Myo1C* but is still weakly retained to an extent sufficient to neutralize the right-handed activity of wild-type macrophages. Altogether, the four loop regions of Myo1C retain at least some of the left-handed information that is functional in the chirality of the hindgut and intracellular F-actin dynamics.

**Fig. 4.**
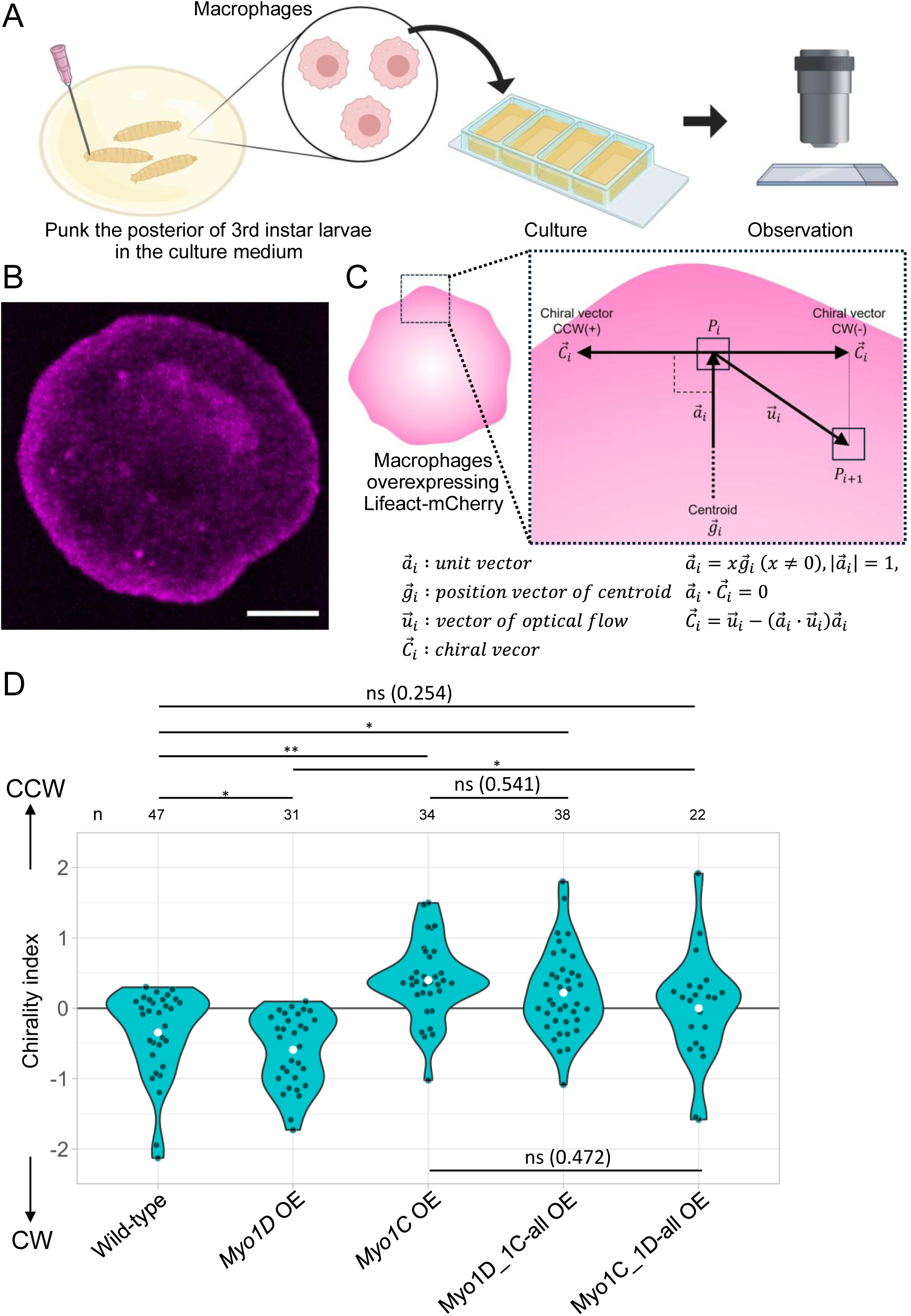
Loop swapping affects the chirality of intracellular F-actin flow in *Drosophila* macrophages. **(A)** Schematic diagram summarizing the *ex vivo* macrophage culture procedure. **(B)** A snapshot forms a live-imaging movie of a wild-type macrophage expressing *Lifeact-mCherry* (magenta). A scale bar represents 5 μm. **(C)** Schematic illustration of chiral vector calculation. The magnified right panel highlights a macrophage expressing *Lifeact-mCherry*; each box represents a pixel at time points i and i+1. F-actin flow was estimated by the optical flow analysis of each pixel, which is defined as a “vector of optical flow.” The direction vector of the optical flow was divided into two vectors to reveal the chirality of F-actin flow: the position vector of the centroid (toward the cell centroid) and the “chiral vector” (perpendicular to the position vector of the centroid). The chirality of F-actin flow was defined by the chiral vector with CW (−) or CCW (+) rotational directionality. **(D)** Violin plot showing F-actin flow chirality index. Each gray dot marks an index obtained from an individual macrophage. Negative and positive values indicate clockwise and counterclockwise rotation biases, respectively. White dots indicate mean values. The number of macrophages analyzed (“n”) is shown above the graph. *P*-values were obtained by Steel–Dwass; ns, *, and ** correspond to *P* > 0.05, *P* < 0.05, and *P* < 0.0001, respectively.

**Table 1.**
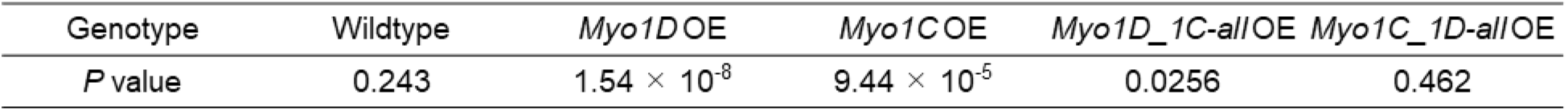
Statistical significance of CW or CCW biases in the chiral F-actin flow. The *P* values of the statistical significance of CW or CCW biases in the chirality index of the F-actin flow in the macrophages of the indicated genotypes were calculated via a one-sample one-sided Wilcoxon signed-rank test.

### Swapping the loop regions changes the interaction with F-actin predicted by the molecular dynamics simulation

We found that the loop regions of Myo1C have a specific function in inducing left-handed activity in the hindgut and that the chiral F-actin flow mediates this function.

Thus, to understand what molecular characteristics of the interaction between the loop regions and F-actin, we performed the molecular dynamics (MD) simulations and calculated the binding free energy of F-actin with Myo1D, with Myo1C, and with their swapped derivatives by the molecular mechanics Poisson-Boltzmann surface area (MM- PBSA) method(*54*) (Figure 5A to 5G, and S4). Briefly, the binding free energy is calculated as the difference between the F-actin–Myosin complex and the individual molecules, each in implicit solvent. The more minor (larger negative) binding free energy means more stable and stronger interactions in the complex. Using AlphaFold3, we built the protein complexes of the three actin monomers, each with ADP and Mg^2+^ in the binding pocket, and wild-type Myo1D, wild-type Myo1C, Myo1D_1Cloop-all, and Myo1C_1Dloop-all. After solvating the protein complexes in explicit solvent and performing equilibration steps, we ran five independent 500 ns-long MD simulations for each system. The contributions of the loops to the total binding free energy were calculated using MM-PBSA with per-residue decomposition (Figures 5A-5C). We detected negative values of MM-PBSA between actin monomer and loop2, loop3, and cm loop, but not loop4, in wild-type Myo1D, wild-type Myo1C, Myo1D_1Cloop-all, and Myo1C_1Dloop-all. This indicates that loop2, loop3, and cm loop, but not loop4, stably bind to actin (Figures 5A to 5C). The inability to detect the binding between loop4 and actin is consistent with the structures predicted by AlphaFold3, which indicate that loop4 is located far from the interface with actin (Figure 1A). Although loop2, loop3, and cm loop have similar total binding free energies, our loop swapping analyses of Myo1D and Myo1C *in vivo* showed that loop2 and loop3 have more notable effect than cm loop on dictating the right- or left-handed rotations of the hindgut (Figure 2F -④, ⑤, ⑩ and ⑪), suggesting that the difference in the binding energies of loop2 and loop3 between Myo1D and Myo1C defines right- or left-handed chirality. We found that the mean total binding free energy in loop2 and loop3 of wild-type Myo1C was smaller than that of the wild-type Myo1D (Figure 5A). In addition, loop3 of wild-type Myo1C has a larger standard deviation than that of wild-type Myo1D and other loops of Myo1D and Myo1C, suggesting that loop3 of Myo1C is flexible and has multiple conformations to interact with actin (Figure 5A).

**Fig. 5.**
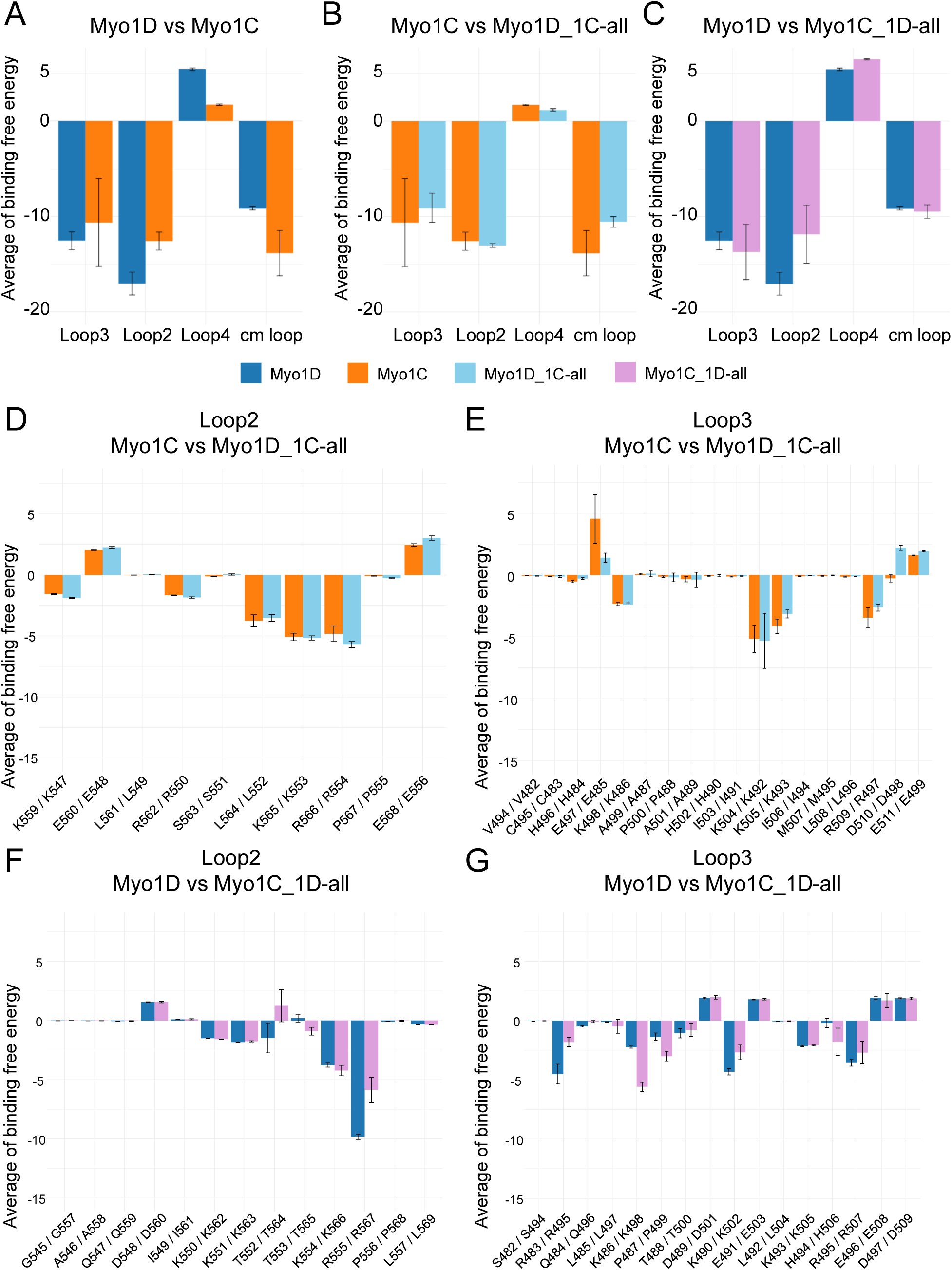
The average values of binding free energy between each amino acid in the loop regions and actin calculated by MM-PBSA analysis and MD simulations. Comparison of the average value of binding free energy between Myo1D and Myo1C **(A)**, Myo1C and Myo1D_1Cloop-all **(B)**, Myo1D and Myo1C_1Dloop-all **(C)**. Blue, Myo1D; orange, Myo1C; light blue, Myo1D_1Cloop-all; purple, Myo1C_1Dloop-all. **(D, E)** The average binding free energy in the loop2 (D) and loop3 (E) is compared between Myo1C and Myo1D_1Cloop-all. **(F, G)** The average binding free energy in the loop2 (F) and loop3 (G) is compared between Myo1D and Myo1C_1Dloop-all (**G**).

MM-PBSA predicted that loop2 of Myo1C interacts with actin through the amino acids of L564, K565, and R566 (Figure 5D). Notably, loop2 of Myo1D_1Cloop-all showed almost the same binding free energy at the corresponding amino acids in the loop2 of wild-type Myo1C (Figure 5D). Similarly, loop3 of wild-type Myo1C interacts with actin through amino acids K504, K505, and R509, and loop3 of Myo1D_1Cloop-all showed similar profiles of binding free energy at the corresponding amino acids in loop3 of wild-type Myo1C (Figure 5E). Moreover, cm loop of wild-type Myo1C interacts with actin through amino acid R319, and cm loop of Myo1D_1Cloop-all showed similar profiles of binding free energy at the corresponding amino acids in the cm loop of wild-type Myo1C (Figure S4A). These results suggest that the profiles of the binding free energy associated with the amino acids in loop2, loop3, and cm loop of Myo1C are maintained even after transferring to the backbone of Myo1D. This result is consistent with the fact that swapping the four loops of Myo1D with the corresponding loops of Myo1C causes a change in chirality-inducing activity from right- to left-handed in the swapped Myo1D.

On the other hand, loop2 of wild-type Myo1D interacts with actin through the amino acids of K554 and R555 (Figure 5F). Interestingly, Myo1C_1Dloop-all showed weaker interactions with actin through amino acids T564 and R567 that correspond to the T552 and R555 of Myo1D, respectively (Figure 5F). Moreover, loop3 of wild-type Myo1D interacts with actin through the amino acids of R483, K490, and R495 (Figure 5G). We found that Myo1C_1Dloop-all did not show the same interaction with actin through the amino acid of R495 that corresponds to R483 of Myo1D, but showed stronger interaction through K498 (which is K486 in wild-type Myo1D) (Figure 5G). Therefore, the profiles of the binding free energy associated with the amino acids in loop2 and loop3 of Myo1D are not maintained when loop2, loop3, loop4, and cm loop of Myo1C were substituted for the corresponding loops of Myo1D in the Myo1C backbone. Such disruption of interactions between the four loop motifs and actin may explain our results, which show that simultaneously replacing the four loops of Myo1D with the Myo1C backbone eliminates both right- and left-handed activity *in vivo*.

## Discussion

In this study, we analyzed the roles of the loop regions of *Drosophila* Myo1D and Myo1C in determining whether cells and organs adopt left- or right-handed chirality. Here, we here found that the four loop regions — loop 2, loop 3, loop 4, and cm loop — of Myo1C together contain sufficient information to confer left-handed chirality, revealing a molecular basis for left-right asymmetry. Although several proteins have been implicated in determining cellular or organ chirality, our work identifies specific motifs that determine whether a cell or organ adopts left- or right-handed chirality. For example, structural studies of the basal body, which controls the clockwise rotation of nodal cilia in mice(*9*), have suggested that it serves as the origin of body chirality, yet the region of the protein responsible for determining the direction of rotation remains poorly understood. Likewise, while the rotational direction of Formin-mediated actin polymerization has been linked to cytoplasmic chirality(*25*), the protein motif by which Formin rotates F-actin is not well understood, nor is it clear how this mechanism relates to cellular chirality.

To dissect the molecular basis of Myo1D and Myo1C activities in determining whether to adopt left- or right-handed chirality, we exchanged their four-loop regions and examined whether cellular and organ chirality switched accordingly. We revealed that the left-handed chirality-inducing activity of Myo1C loops emerged only when all four loops were transferred together to the Myo1D backbone, suggesting that cooperative interactions and spatial coordination among these loops are required. MM-PBSA analysis further revealed that Myo1C loops maintained similar actin-binding profiles even when grafted onto the Myo1D backbone, indicating a degree of structural autonomy when placed in three-dimensional space. This may explain why transferring all four Myo1C loops to the Myo1D backbone conferred left-handed activity. In contrast, Myo1D loops transplanted into the Myo1C backbone displayed altered actin-binding profiles compared to those of wild-type Myo1D, implying that Myo1D loops function depends more strongly on the surrounding structural context. Consequently, the right-handed activity of Myo1D loops might not be fully captured in this chimeric system. According to this view, it cannot be ruled out that the four loops of Myo1D contain sufficient information that confers the right-handed chirality in Myo1D’s backbone. However, the swapping approach implemented in this study may induce nonspecific disruption of protein structure by exchanging loop regions, potentially inhibiting activity in determining whether to adopt left- or right-handed chirality.

Therefore, if the chirality-inducing activities of the swapped Myo1C and Myo1D decrease, this cannot be used as evidence to conclude that the swapped loop regions possess sufficient information for inducing left- or right-handed chirality.

Although the possibility of protein denaturation due to loop-region swapping cannot be ruled out, our analysis using the embryos pinpointed loop2 and loop3, but not loop4 or cm loop, as critical regions for the right- and left-handed activities of Myo1D and Myo1C, respectively (Figure 2F). Moreover, our MM-PBSA analysis revealed that loops 2 and 3 of Myo1D bind actin more tightly than those of Myo1C, a difference that may underlie their opposing chiral outcomes. Based on these results, we hypothesize that the difference in binding strength between loop2 and loop3 and actin determines whether left- or right-handed chirality is induced. However, further analysis considering protein structure is required to evaluate this possibility.

Previous biochemical assays have shown that Myo1D and Myo1C differ in how they move F-actin with respect to chirality(*35, 37*). In a standard *in vitro* motility assay used in the previous studies, Myo1D moves F-actin in a clockwise direction, whereas Myo1C drives it linearly or randomly(*35, 37*). At higher actin concentrations used in this study, Myo1D bundles F-actins into clockwise-rotating rings, while Myo1C induces non-directional flow(*51*). In addition, recent time-resolved cryo-electron microscopy studies of mouse myosin V have captured the initial weak actin-binding state and revealed the ordered sequence of events underlying force production(*55*). According to this model, myosin initially engages with actin via loops 2 and 3, triggering subsequent rearrangements of the head subdomains, phosphate release, and the execution of the lever-arm power stroke. During this sequence, the upper 50-kDa subdomain of the myosin head, which includes loop2, cm loop, and loop4, undergoes a clockwise rotation around the actin filament axis when viewed from the myosin perspective and binds the actin strongly through cm loop^55^. We speculate that this rotational movement contributes to the directional bias of actin filament motility. During the transition to strong actin binding, binding through loop 3 may stabilize the clockwise rotation of the upper 50-kDa subdomain, thereby facilitating filament-actin’s LR-directional bending. Our MD simulation suggests that the average binding free energy in loop3 of Myo1D is higher and more stable than that of Myo1C (Figure 5A). Thus, we speculate that the strong binding to actin through loop3 stabilizes the clockwise rotation of the upper 50-kDa subdomain in Myo1D and enables the bending motion of F-actin, while the rotation of the upper 50 kDa is not supported in Myo1C because of unstable binding between loop3 and actin, leading to the low bending of F-actin’s motion. In agreement with this hypothesis, swapping loop 3 between Myo1D and Myo1C reduced their respective right- and left-handed activities in the LR-asymmetric rotation of the embryonic hindgut (Figure 2F -row⑤ and ⑪).

Comparative structural analyses also indicate that loop2 footprints on actin are highly conserved across myosins, while those of loop3, loop4, and cm loop are more divergent, likely reflecting functional specialization(*56*). These observations also support our idea that loop3 plays specific roles in determining cell and organ chirality.

Together, our findings reveal that the four loop regions of Myo1C contain molecular information sufficient to determine whether cells and organs adopt left- or right-handed chirality. However, the molecular mechanism by which the Myo1C loop motifs drive counterclockwise F-actin spiraling in macrophages remains unknown. Specifically, in the *in vitro* motility assay containing a high concentration of F-actin, Myo1D bundles F-actin into clockwise-rotating rings, whereas Myo1C induces only a random flow of F-actin(*35, 37, 40*). Therefore, the origin of the chirality of the force that causes F-actin to spiral in a left-handed manner, as seen in wild-type Myo1C and Myo1D_1Cloop-all in macrophages, remains unknown. Our genetic epistasis analysis revealed that even in the absence of *Myo1D* function, *Myo1C* retains left-handed activity independently of *Myo1D* (Figure 2F -row③). Therefore, Myo1D and Myo1C may induce right- and left-handed chirality, respectively, through distinct mechanisms. This finding implies that multiple molecular systems operate in parallel to establish LR asymmetry in *Drosophila*. Nevertheless, together, our results provide a mechanistic framework linking myosin–actin interactions to the emergence of cellular and organismal chirality.

## Materials and Methods

### Generation of plasmids and transgenic flies

To construct pUASt-Myo1C-mCherry, PCR fragments of *Myo1C* and *mCherry* cDNAs were combined by PCR, and the resulting fragment was cloned into the pUASt vector, using the same methods previously described(*51*). To construct pUASt-Myo1D-mGFP6, PCR was used to combine PCR fragments of Myo1D and mGFP6 cDNAs, and the resulting fragment was cloned into the pUASt vector(*43, 51*).

To generate loop-region-swapped Myo1D and Myo1C genes, we replaced the loop-region cDNAs in the recipient myosins with the corresponding sequences from the donor myosins using PCR. Briefly, using Myo1D or Myo1C cDNA as template, PCR fragments were generated by primers designed to introduce the donor myosin-derived sequence at the 3′ end of one fragment and at the 5′ end of the other. These fragments share the donor myosin-derived overlaps and were fused into a single chimeric fragment by overlap extension PCR. The resulting products were cloned into the corresponding sites of pUASt-Myo1D-mGFP6 or pUASt-Myo1C-mCherry. pUASt-Myo1D_1Cloop-2 and pUASt-Myo1C_1Dloop-2 were subsequently used as the template to replace the additional regions of loop3 of recipient myosins with the corresponding sequences from donor myosins by the same method. The procedure was repeated sequentially to introduce multiple donor myosin-derived loop regions into the construct, yielding pUASt-Myo1D_1Cloop-all and pUASt-Myo1C_1Dloop-all. For all loop-region-swapped Myo1D and Myo1C genes, their nucleotide sequences were confirmed by DNA sequencing. These constructs were integrated into the 68A4 P[CaryP]attP2 site of the third chromosome by using the PhiC31/attP/attB system to obtain the transgenic *Drosophila* lines(*57*).

### Fly lines

All genetic crosses were performed on a standard *Drosophila* culture medium at 25 °C. We used *white^1118^* (*w^1118^*), carrying a *w* mutant otherwise wild type, as the wild-type strain. For *Myo1D* null alleles, we used *Myo1D^L152^*strains(*32*). The following GAL4 lines were used in this study: *byn*-GAL4, a hindgut-specific driver(*50*), and *He*-GAL4, a macrophage-specific driver (Bloomington #8699). We used *UAS-Lifeact-mGFP*6 and *UAS-Lifeact-mCherry* for labeling F-actin(*51*).

### Staining of embryos

Antibody staining was carried out as previously described(*32, 36*). The following primary antibodies were used at the dilution indicated: rabbit anti-RFP (1:1,000, MBL), rabbit anti-GFP (1:1000, MBL), mouse anti-Fasciclin III (1:100, 7G10, Developmental Studies Hybridoma Bank), and chicken anti-β-galactosidase (1:500, Abcam).

The following secondary antibodies were used at the dilution indicated: anti-rabbit IgG-Cy3 (1:500, Jackson ImmunoResearch), anti-rabbit IgG-Alexa Fluor 488 (1:500, Molecular Probes), anti-mouse IgG-Cy3 (1:500, Jackson ImmunoResearch), anti-mouse IgG-Alexa Fluor 488 (1:500, Molecular Probes), anti-chicken IgY-Alexa 488 (1:500, Jackson ImmunoResearch).

### Microscopic analysis of embryonic gut handedness

*Myo1D^L152^* homozygous embryos were identified by the absence of blue balancers, as indicated by the lack of β-galactosidase staining. To evaluate LR asymmetry of the hindgut, embryos were mounted in 80% glycerol and observed with a differential interference microscope (Axioskop 2 plus, Carl Zeiss). The images were obtained by confocal microscopy (LSM880, Carl Zeiss).

### Preparation of macrophages for live imaging and Optical flow analysis of F-actin

*Ex vivo* culture of macrophages was carried out as previously described(*51*). Briefly, the macrophages were collected from 3^rd^-instar larvae by poking the posterior body in M3 medium (Sigma). The medium containing the macrophages was transferred to a chamber dish (Thermo Fisher) coated with Concanavalin A (Nacalai Tesque) and cultured at 27 °C for 30 min. Live images were obtained every 15 seconds for 10 minutes using an LSM 880 (Carl Zeiss) and processed with Airyscan (Carl Zeiss).

These acquired time-lapse images were processed for the optical flow analysis as previously described(*51*). Briefly, the optical flow vector for all pixels was estimated using the OpenCV optical flow analysis with Gunnar Farnebäck’s algorithm(*53*). The chiral vector and chirality index of F-actin flow were calculated as described in the results section.

### Western blot analysis

Each protein sample was prepared from thirty embryos. The embryos were homogenized in RIPA buffer (150 mM NaCl, 50 mM Tris-HCl (pH 8.0), 1% Triton X-100) with Protease Inhibitor (Sigma-Aldrich). Homogenized embryos were sonicated and centrifuged for 5 min at 15000 rpm. The supernatants were incubated for 3 min at 95°C, and the protein extracts were mixed in the Sample buffer. The protein extracts were loaded into 7 % acrylamide SDS-PAGE gel. The separated proteins were transferred onto PVDF (Poly Vinyli dene Fluoride). The following primary antibodies were used at the dilution indicated: rabbit anti-RFP (1:5000, MBL), rabbit anti-GFP (1:1000, MBL), and mouse anti-alpha-tubulin (1:5000, NOVUS BIOLOGICALS). As the second antibodies, ECL peroxidase labelled anti-mouse antibody (1:5000, Cytiva), HRP-conjugated anti-rabbit antibody (1:5000, Proteintech) were used. To detect the signals, SuperSignal West Femto Maximum Sensitivity Substrate was used (Thermos Scientific).

### Statistical analysis

Data were statistically analyzed using R (version 3. 6. 1). Fisher’s exact test with Holm correction was performed (Fig. 2F). P-values were calculated via Steel–Dwass (Fig. 4D). One-sample one-sided Wilcoxon signed rank test was performed (Tables 1).

### Prediction of protein structures by AlphaFold3

Protein structure prediction was performed using AlphaFold3(*47*). The amino acid sequences of myosins and actin (Uniprot: Q23978 (Myo1D), Q23979 (Myo1C), P10987 (Actin 5C)) were used as input, including 3 ADP molecules and 3 Mg^2+^ ions.

Predictions were run with default parameters on Google Colab using the AlphaFold Server, and the best-ranked models, as determined by the pLDDT confidence score, were selected. Predicted structures were visualized using Chimera X v1.9.

### Molecular dynamics simulation and MM-PBSA analysis

The AlphaFold 3-generated complex structures were used as initial structures for MD simulations. Each structure, which consists of one myosin and three actin molecules with an ADP and a magnesium ion in each binding pocket, was neutralized using sodium ions and solvated in a water box with ∼78,000 water molecules. Amber ff14SB force field(*58*)was used for the protein, ADP was treated with the force field by Meagher et al.(*59*), and TIP3P water model was adopted. Long-range electrostatic interactions were calculated using the particle mesh Ewald method(*60*), and short-range non-bonded interactions were cut off at 10.0 Å.

Each system was energy-minimized in three steps: 3,000 steps with heavy-atom restraints, 3,000 steps with backbone-atom restraints (N, CA, C, O), and 3,000 steps without restraints. Each system was then heated up to 300 K in 50 ps, followed by a constant-NPT (300 K, 1 atm) simulation for 500 ps while restraining the protein with a weak force constant of 1 kcal mol^-1^ Å^-2^ and another 500 ps with a restraint on the backbone atoms of actin with a force constant of 1 kcal mol^-1^ Å^-2^. Finally, another 20 ns simulation under constant-NVT (300 K) with the same restraint on the backbone atoms of actin was performed to complete the equilibration steps, while collecting structures every 4 ns for subsequent production runs. From the five structures collected during equilibration, production simulations were performed for 500 ns under the constant-NVT (300 K) condition with a 1 kcal mol^-1^ Å^-2^ restraint on the backbone atoms of actin using different random seeds. The coordinates were recorded every 1 ns.

The interaction energies between myosin and actin (including ADP molecules and magnesium ions) were calculated with the molecular mechanics Poisson-Boltzmann surface area (MM-PBSA) method after extracting the water molecules and sodium ions from the trajectory. The dielectric constants in the protein and water were set to 1 and 80, respectively. Energy decomposition was performed on a per-residue basis, and the binding energy contributions from the loops were calculated by summing over the residues in the loops. The mean values and error bars were estimated from the mean and standard deviation of the five trajectories calculated per system. All MD simulations and MM-PBSA calculations were performed using the Amber 20 package(*61, 62*)and the MMPBSA.py program in Amber(*54*), respectively.

## Acknowledgements

We thank the members of the Matsuno Laboratory for their valuable advice and discussions. We also thank the Bloomington *Drosophila* Stock Center (Indiana University) and the Kyoto Stock Center (Kyoto Institute of Technology) for *Drosophila* stocks. This work was supported by the Grant-in-Aid for Transformative Research Areas (15H05863 and 25K09646) to K.M.; and by the Grant-in-Aid for JSPS Fellows (22J20544) to A.Y.

## Author contributions

K.M. and K.I. conceived and directed the study. A.Y. and T.S. performed the observation of embryos and macrophages. A.Y. and T.S. performed the data analysis. K. Y., T. H. supervised the experiments. T. M. conducted MD simulations and MM-PBSA calculations. K.M., K.I., A.Y., and T. M. wrote the draft. All authors approved the final version of the manuscript.

## Lead contact

Requests for further information and resources should be directed to and will be fulfilled by the lead contact, Kenji Matsuno

## Data and materials availability

Any additional information required to reanalyze the data reported in this paper is available from the lead contact upon request.

---

**Fig. S1.**
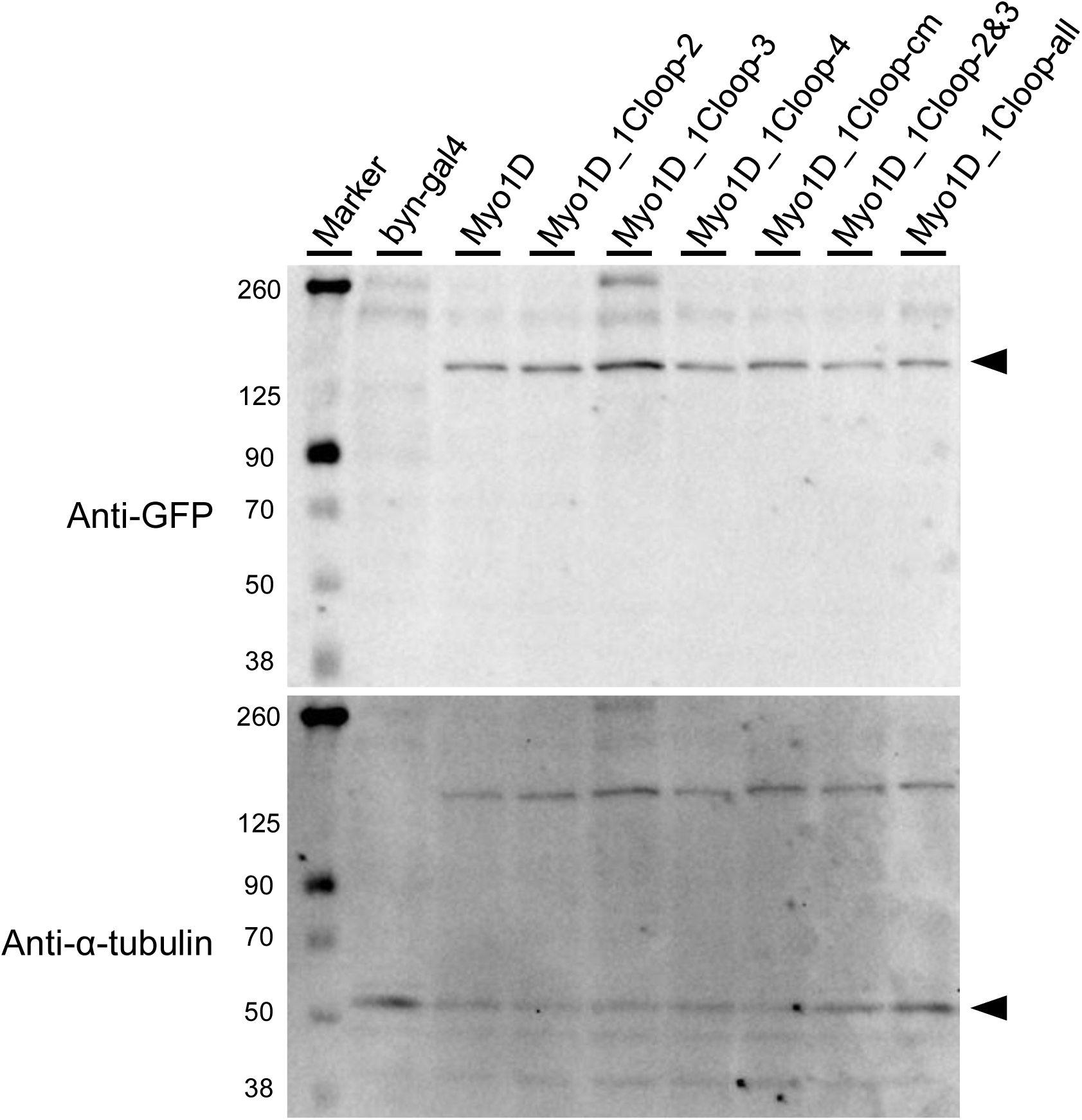
The wild-type and chimeric Myo1D derivatives were properly detected on Western blot analysis. (Top) Myo1D and the mGFP6-tagged chimeric Myo1D were detected at approximately 144 kDa using an anti-GFP antibody. The protein extracts were obtained from wild-type embryos overexpressing corresponding genes in the hindgut. (Bottom) α-tubulin for each sample was detected as a loading control (55 kDa) after stripping the same membrane shown at the top. Molecular weight markers are presented at the left of each blot. *byn-gal4* was used for the negative control.

**Fig. S2.**
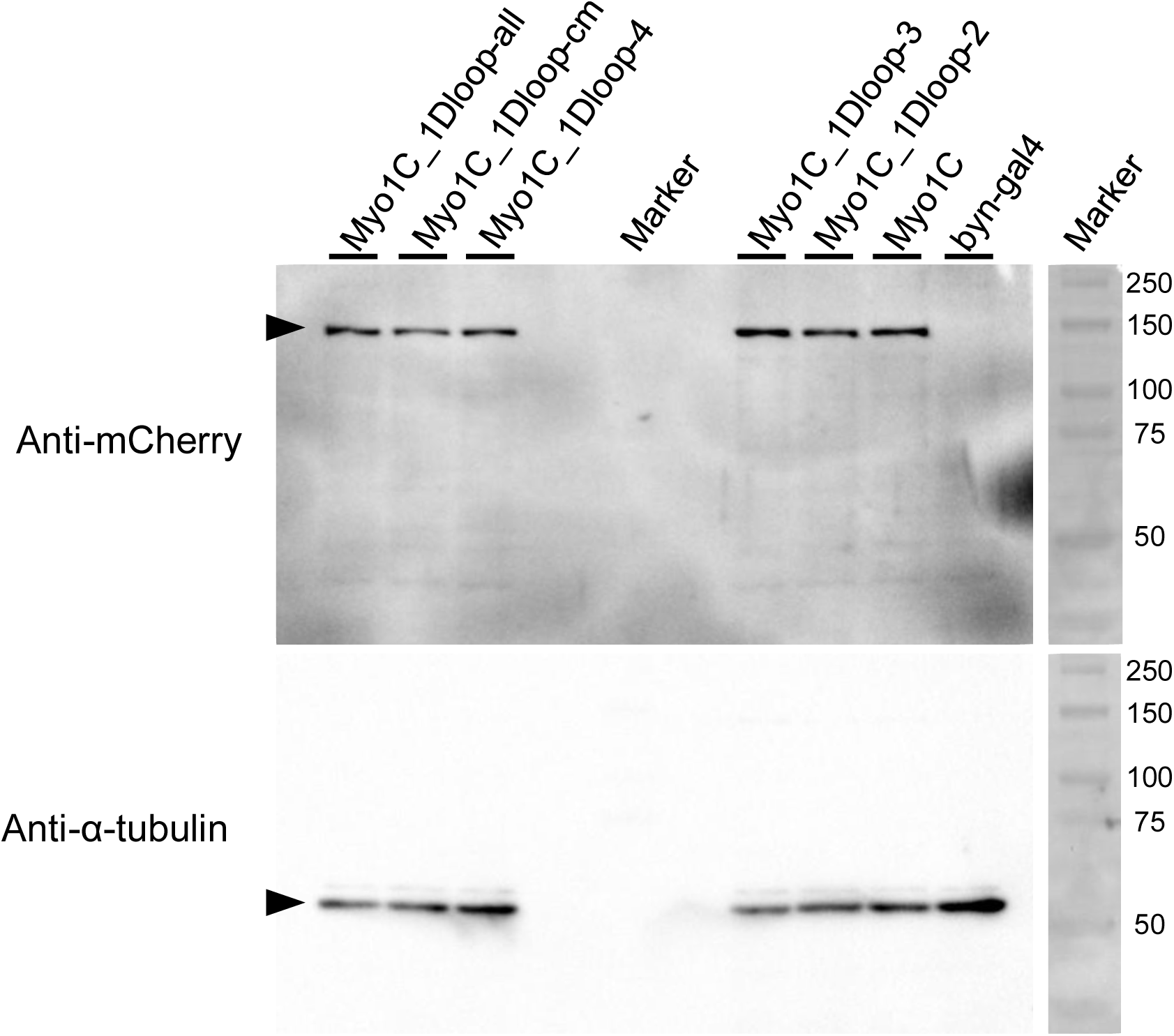
The wild-type and chimeric Myo1C derivatives were properly detected on Western blot analysis. (Top Left) Myo1C and the chimeric Myo1C tagged with mCherry were detected at the position of around 144 kDa using an anti-RFP antibody. The protein extracts were obtained from wild-type embryos overexpressing corresponding genes in the hindgut. (Bottom left) α-tubulin in each sample was detected as a loading control (55 kDa) after stripping the same membrane shown at the top. (Top and bottom right) Molecular weight markers. *byn-gal4* was used for the negative control.

**Fig. S3.**
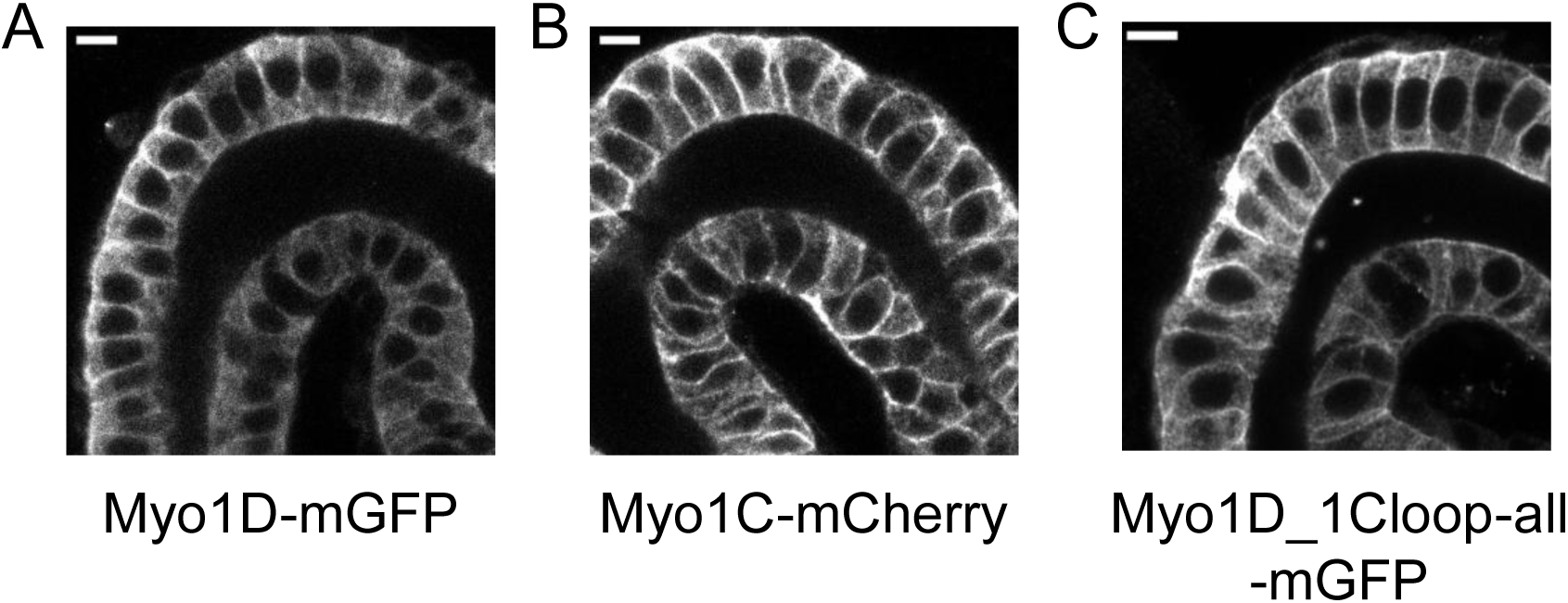
No differences among Myo1D-mGFP6, Myo1C-mCherry and Myo1D_1Cloop-all-mCherry were detected in the intracellular localization of the three proteins in the embryonic hindgut epithelium. Localizations of Myo1D-mGFP6 (**A**), Myo1C-mCherry (**B**), and Myo1D_1Cloop-all-mCherry (**C**) were shown. The pictures show the hindgut portion in the fixed embryos. Each myosin was stained with antibodies specific to GFP or mCherry.

**Fig S4.**
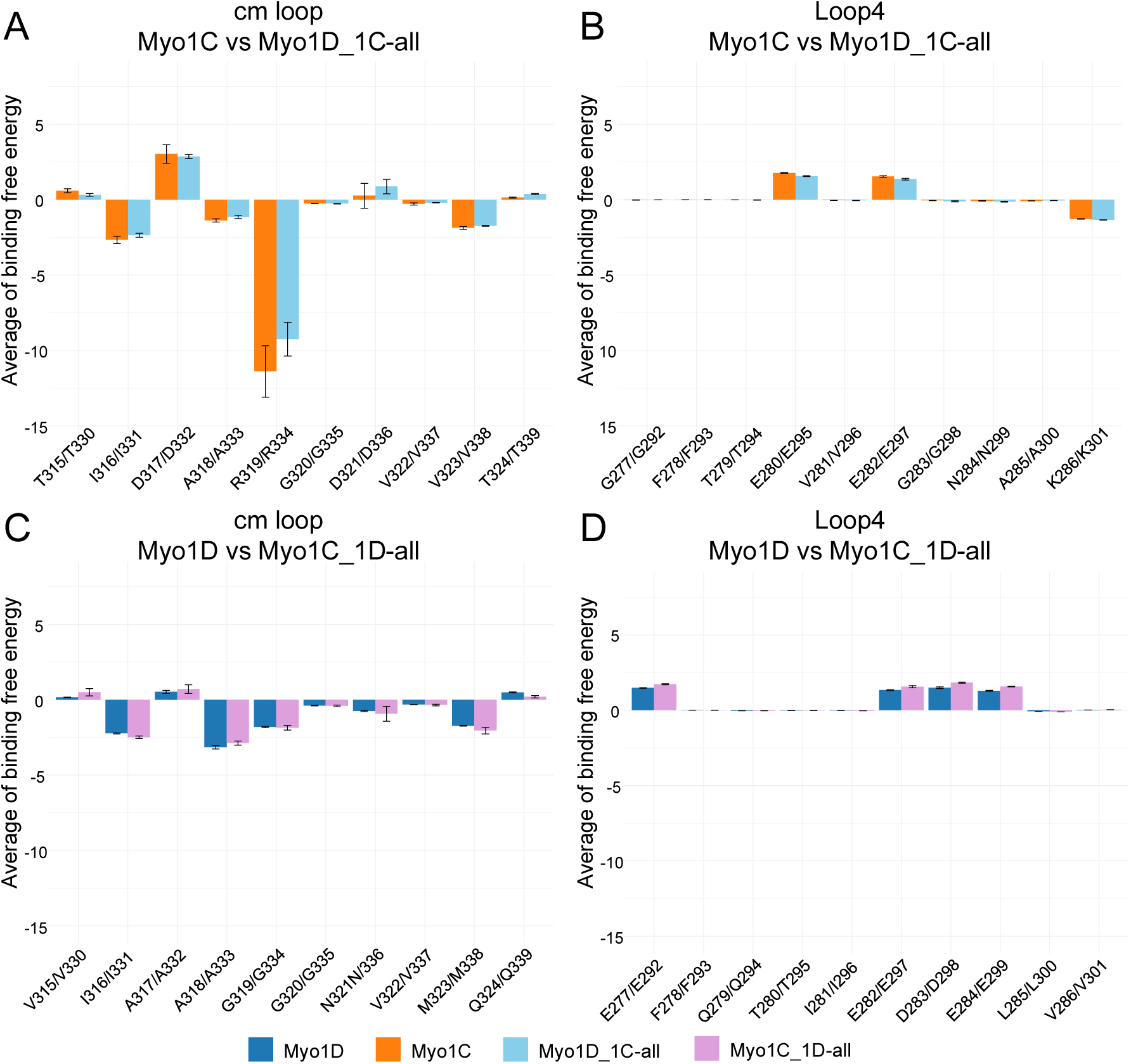
The average binding free energy between each amino acid in the loop regions and actin was calculated using MM-PBSA analysis and MD simulations. (**A**, **C**) For the cm loop, panel A compares Myo1C and Myo1D_1Cloop-all, while panel C compares Myo1D and Myo1C_1Dloop-all. (**B**, **D**) For loop4, panel B compares Myo1C and Myo1D_1Cloop-all, while panel **D** compares Myo1D and Myo1C_1Dloop-all. Blue, Myo1D; orange, Myo1C; light blue, Myo1D_1Cloop-all; and purple, Myo1C_1Dloop-all.

**Supplementary Movie 1**

A time-lapse movie of a wild-type embryonic hindgut expressing *Lifeact-mGFP* for 2 h: Lifeact-mGFP is shown in green. Scale bar is 20 μm.

**Supplementary Movie 2**

Time-lapse movies of macrophages overexpressing *Lifeact-mGFP6 or Lifeact-mCherry*; wild type, *Myo1D* overexpression (OE), *Myo1C* OE, *Myo1D_1Cloop-all* OE, *Myo1C_1Dloop-all* OE for 10 min. Lifeact-mGFP6 and Lifeact-mCherry are shown in white. Scale bars are 5 μm.

